# CRISPRware: an efficient method for contextual gRNA library design

**DOI:** 10.1101/2024.06.18.599405

**Authors:** Eric Malekos, Christy Montano, Susan Carpenter

## Abstract

We present CRISPRware, an efficient method for generating guide RNA (gRNA) libraries against transcribed, translated, and noncoding regions. CRISPRware leverages next-generation sequencing data to design context-specific gRNAs and accounts for genetic variation, which allows allele-specific guide design on a genome-wide scale. The latter ability holds promise for the development of gene therapy in the context of gene dosing and dominant negative mutations.

## Main Text

Screening methods are useful for uncovering associations between genomic elements and molecular pathways or cellular phenotypes. CRISPR screens have become a popular screening modality due to CRISPR-Cas9’s ability to efficiently knockout protein-coding genes with high specificity. Moreover, CRISPR interference (CRISPRi) and CRISPR activation (CRISPRa) systems allow transcriptional repression or overexpression as well as modulation of cis-regulatory elements. Contemporaneously, the molecular biology community has embraced next-generation sequencing (NGS) approaches, generating datasets that can greatly enhance CRISPR guide RNA (gRNA) design by describing the genomic and transcriptomic landscapes in a given experimental context^1,2^. Yet, despite the importance of gRNA library design for successful CRISPR experiments, there has not been a computational tool with native support for contextual NGS data.

CRISPRware is a Python software package that can use various processed NGS data types to generate gRNA libraries. The workflow starts by defining genomic targets of interest (e.g. gene coding sequences or transcription factor binding sites), determining available protospacers in target regions, scoring the putative protospacer-targeting gRNAs for on-target and off-target activity, and returning a ranked list of gRNAs for each target (Fig 1A). Off-target activity is determined by GuideScan2, a well-validated tool for this purpose^3–5^. On-target scoring (or cleavage efficiency) is determined by Ruleset 3^6^. To demonstrate the core functionality of CRISPRware, we document all available NGG (Cas9) and TTTV (Cas12A or Cpf1) protospacers in the coding sequences (CDS) of NCBI RefSeq genes (Fig 1B, Supp Fig 1A, Supp Tables 1-12). On-target scoring has been shown to benefit from an ensemble approach combining multiple scoring methods^7^. Therefore, we made CRISPRware compatible with many other on-target scoring methods via integration with the crisprVerse Bioconductor package^8^(Fig 1C). To facilitate the widespread use of CRISPRware, we implement a batch-size parameter that allows users to score gRNAs efficiently even on personal computers with limited resources (Supp Fig 1B).

**Figure 1.**
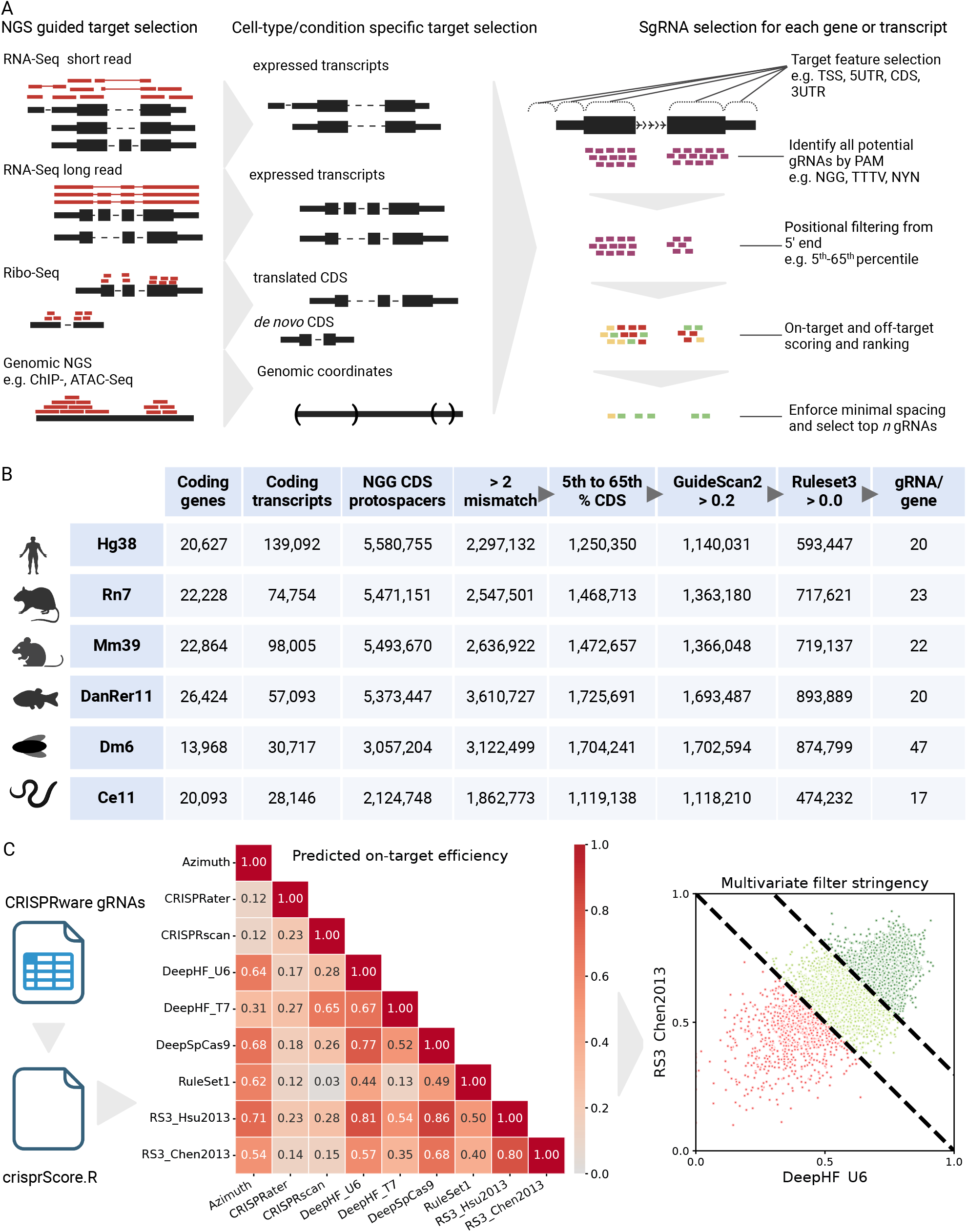
**(A)** Overview of CRISPRware gRNA selection pipeline. Left, CRISPRware takes a genomic fasta and a target file (GTF/GFF or BED). Left and center, optionally, a variety of processed NGS data can be used to inform target selection. Right, CRISPRware implements methods for on-target and off-target scoring and established practices for selecting a top set of gRNAs for genes and transcripts. **(B)** Demonstration of CRISPRware applied on model genomes with NCBI RefSeq gene annotations. After identifying protospacers (column 3), the standard CRISPRware filtering pipeline is applied, with subsequent columns applying all previous filters. 5th-65th percentile CDS filtering is based on previous reporting^10^. The final column reports the median gRNAs available for each gene, one of the summary readouts provided at each stage of filtering. **(C)** Left, CRISPRware is packaged with Ruleset 3 (RS3) but has built-in interoperability with the crisprScore module of crisprVerse, which allows many additional scoring methods^6,10,20,21,21–23^. Center, Spearman correlation matrix of crisprScore Cas9 techniques targeting Hg38 coding sequences. Right, RS3 scoring with Chen2013 tracr^6^ versus DeepHF^23^ scoring with U6 promoter after applying max-min normalization to both methods. 5,000 randomly selected Hg38 CDS-targeting guides are displayed.

While the number of annotated protein-coding genes has stabilized, novel protein-coding isoforms, transcription start sites, noncoding genes, and genes that encode small peptides continue to be discovered (Sup Fig 2A-D)^9^. The isoform expression, TSS usage, and translation of noncanonical reading frames are cell-type and context-specific. These observations argue against the use of static CRISPR gRNA libraries^10^ and towards contextual library design. For instance, CRISPRware can make use of RNA-Seq transcript per million (TPM) data from popular long-read^11,12^ and short-read tools^13,14^ to filter based on isoform expression. Depending on the desired complexity of the gRNA library, the user can target all isoforms above some defined TPM cut-off or generate new gene models with a single isoform-per-gene (Fig 2A). In testing human cell lines and mouse leukocyte cells at various stringency cut-offs, we found that a consensus gene model, composed of shared exons and CDS features of all isoform variants for a given gene, could almost always be constructed (Sup Fig 2 E-F). Using such models solves the optimization problem of choosing the fewest gRNAs with the highest likelihood of knocking out gene function.

**Figure 2.**
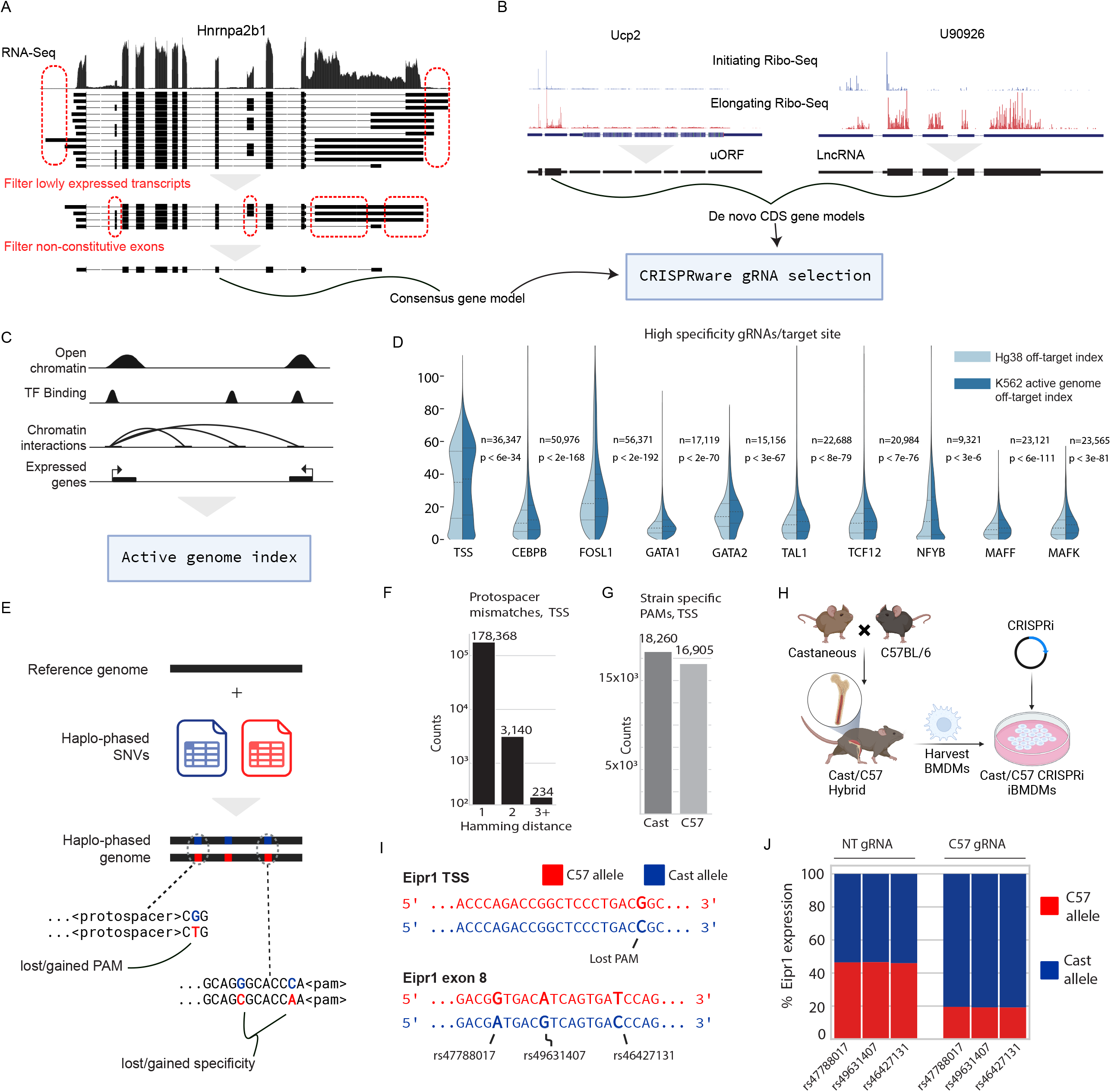
**(A)** Demonstration of RNA-Seq guided target selection. Top, RNA-Seq at Hnrnpa2b1 locus in from mouse bone marrow-derived macrophages (BMDMs). Middle, remaining transcripts after expression filtering. Bottom, consensus gene models are constructed from shared isoform regions. **(B)** Demonstration of Ribo-Seq guided target selection. *De novo* gene models are built from noncoding regions with predicted translated open reading frames **(C)** Schematic of CRISPRware cell type-specific off-target database construction from processed genomic NGS data. **(D)** Availability of gRNAs with high specificity (GuideScan2 score > 0.2) against either the entire Hg38 genome or the active portions of the genome in K562. See Methods for construction of K562 active genome. **(E)** Schematic of CRISPRware allele-specific targeting. **(F**,**G)** Mismatches in the protospacer sequence between C57BL/6 and Castaneous in TSS and PAMs specific to either C57BL/6 or Castaneous appearing in the TSS. TSS-targeting is defined as a +/- 300 bp window around a Gencode vM34 TSS. **(H)** Generation of C57BL/6 x Castaneous CRISPRi BMDMs. **(I)** Top, a PAM site in the TSS of Eipr1 in C57BL/6 is lost in Castaneous. Bottom, three SNPs in exon 8 can be used to discern allele-specific Eipr1 expression. **(J)** Relative expression of Eipr1 from Amplicon-Seq, with either a non-targeting gRNA or a C57 targeting gRNA.

CRISPRware can also make use of processed Ribo-Seq (or Ribosome Footprinting) data to target translated canonical CDSs and novel coding ORFs in regions annotated as noncoding. Recently, CRISPR screen experiments have been used to characterize these novel ORFs and the peptides they encode^15,16^. However, targeting these ORFs is not natively supported by current CRISPR screen design tools. CRISPRware addresses this unmet need by generating *de novo* gene model annotations from Ribo-Seq analysis (Fig 1A, 2B) provided by either of two widely used tools^17,18^, which can be used as input to the CRISPRware pipeline.

CRISPRware supports targeting of noncoding areas such as promoter and enhancer regions for CRISPRi and CRISPRa experiments. Successful targeting of noncoding regions is known to be highly dependent on the chromatin state of the target site^1,2^ and the protospacer position relative to the transcription start site (TSS)^2^. Notably, CRISPRware can combine processed NGS data to leverage these known properties. For example, using RNA-Seq data to determine highly active TSS sites along with either signal tracks (bigwig) or called peaks (BED) from assays against markers of active chromatin (e.g. ATAC-Seq, DNase-Seq, ChIP-Seq, etc.) to target the most accessible window in proximity to the TSS.

One challenge researchers may face in screen design is a lack of suitable gRNAs, especially when adequate off-target filtering is applied^5^. However, based on the observation that CRISPR systems are highly dependent on DNA accessibility^2,19^(PMID: 27353328, PMID: 38504114), we developed a wrapper for GuideScan2 that can index the active portion of the genome (Fig 2C). We demonstrate that the number of targets for which suitable gRNAs can be found for gene TSSs and various transcription factor binding sites is substantially increased for the K562 cell line with this approach (Fig 2D, Supp Fig 3A), even when the active genome is liberally defined (see Methods).

We also accounted for gRNA selection in the context of genetic variation and allele-specific targeting (Fig 2E). We generated a haplo-phased genome by lifting SNPs from the Castaneous (Cast) strain from the Mouse Genome Informatics group onto mouse reference genome Mm39 (C57BL/6 [C57] strain) with a CRISPRware script. CRISPRware was used to identify protospacers with mismatches between the two strains and lost or gained PAM in gene CDSs and TSSs (Fig 2F, G Supp Fig 4A-F). We generated immortalized bone marrow-derived macrophages from a C57-Cast hybrid F1 mouse line and transduced it with CRISPRi machinery (Fig 2H). We used CRISPRware to design guides using RNA-Seq and ATAC-Seq from C57 and knocked down Eipr1. We sequenced the Eipr1 RNA and saw that targeting C57 Eipr1 TSS resulted in reduced expression from the C57 allele, while non-targeting guides did not (Fig 2I, J).

In summary, we present a flexible CRISPR gRNA design tool that can efficiently generate gRNAs at single gene or genome-wide scale. We demonstrate its utility by aggregating CDS-targeting gRNAs scored with multiple methods across six species (Supp tables 1-12). We further demonstrate its ability to generate context-specific gRNA libraries by leveraging publically available NGS data to inform target selection. Finally, we demonstrate CRISPRware’s ability to perform allele-specific targeting, which may facilitate future studies of cis-regulatory interactions, haplo-sufficiency, and dominant negative mutations.

**Supplementary Figure 1 (A)** Demonstration of CRISPRware applied on model genomes with NCBI RefSeq gene annotations, targeting TTTV PAMs. Scores are determined by DeepCpf1^24^ for AsCas12A and enPAMGB^25^ for enCas12a as implemented in crisperVerse^8^. After positional filtering (5th-65th percentile), subsequent filtering is not applied sequentially, and gRNA/gene reports the median available gRNA under each filtering criteria. **(B)** Running enPAMGB scoring on 500,000 gRNAs at various batch sizes. Reducing batch size decreases memory requirements while slightly increasing running time.

**Supplementary Figure 2 (A-D)** Changes in the number of protein-coding genes, protein-coding gene isoforms, and protein-coding isoform TSSs **(A, C)** and from the comprehensive protein-coding and noncoding Gencode releases and **(B, D). (E, F)** The number of expressed genes for which consensus gene models can be constructed at increasingly stringent expression cutoffs.

**Supplementary Figure 3 (A)** Number of target sites, either TSS or transcription factor binding sites, for which there are 0 high specificity gRNAs (GuideScan2 score > 0.2, > 2 mismatches) against either the entire Hg38 genome or the active portions of the genome in K562. See Methods for construction of K562 active genome.

**Supplementary Figure 4 (A-C)** Mismatches in the protospacer sequence between C57BL/6 and Castaneous in CDS regions with NGG pam **(A)**, the TSS with TTTV PAMs **(B)**, and the CDS with TTTV PAMs **(C). (D-F)** Lost/gained PAMs in Castaneous vs. C57BL/6. TSS-targeting is defined as a +/- 300 bp window around a Gencode vM34 TSS.

## Methods

CRISPRware workflow

### RNASeq guided preprocessing

A number of scripts exist to preprocess NGS data prior to determining gRNAs. The module `preprocess_annotation` takes processed RNASeq TPMs from Kallisto/Salmon (short-read) or FLAIR/Mandalorian (long-read) from one or more samples along with the GTF/GFF gene annotation. If multiple samples are passed, max, min, median, and mean TPM values for each transcript are determined, and the user can supply minimum cut-offs for any combination of these to filter out lowly expressed isoforms. All detected isoforms (TPM > 0) are kept by default. The user can also set an integer flag “`--top-n <n>`“ which will filter out all but the `<n>` most highly expressed isoform for each gene.

Following filtering, a filtered GTF is created, along with four optional GTF files: *shortest, longest, metagene*, and *consensus*. Each of these four GTFs will have a single isoform model for a given gene. *Shortest* and *longest* retain the shortest and longest isoform, respectively. *Metagene* constructs a gene model consisting of all exons and coding sequences (CDSs) from all isoforms for a given gene, similar to a union operation. *Consensus* constructs a gene model in which shared or overlapping exons and CDSs from isoforms from a given gene a combined such that exon and CDS entries in the final gene model appear in all other isoforms, similar to an intersection operation. By desgin *shortest, longest*, and *metagene*, generate output gene models for all input genes. *Consensus* models cannot always be generated (e.g. a protein-coding gene in which there are no common CDS entries, Supp Fig 2 E, F), in which case one of the alternative models can be used for those genes.

Optionally, two BED files are generated for each new GTF, a transcription start site (TSS) and a transcription end site (TES) file, which generate coordinates around the TSS and TES in a user-defined window (default: +/- 250 bp).

### RiboSeq guided preprocessing

RiboSeq is a technique that allows researchers to evaluate the coding potential of canonical and noncanonical open reading frames (ORFs) by sequencing ribosome-protected fragments. RiboSeq reads are mapped to a transcriptome, and statistical analyses are performed to infer coding regions. CRISPRware can take ORFs called from RiboSeq and generate new GTFs with *de novo* CDS entries. Currently, GTFs can be generated from the output of either Probabilistic inference of codon activities by an EM algorithm (PRICE) or Ribo TIS Hunter (Ribo-TISH), two widely used RiboSeq processing tools.

### ATACSeq, DNASESeq, ChIPSeq, and other genomic preprocessing

Targeting noncoding elements can be guided by any NGS data that yields BED coordinate files. Additionally, a helper script, `bigwig_to_signalwindow.py`, can take a BED and BigWig signal file and return the window in each BED entry that has the highest mean signal. For example, a TSS BED and ATACSeq BigWig can be paired to determine the area of the TSS that is most nucleosome-depleted - the most important feature for effective CRISPRi experiments.

### Identifying guide RNAs

The first processing module in CRISPware is `generate_guides,` which scans an input genome for protospacers based on a user-defined PAM sequence. GTF/GFF (hereafter GTF) and BED files can be passed to limit the search space to regions of interest. If a GTF is passed, a feature identifier (CDS, 5UTR, 3UTR, exon, etc.) can also be passed to further restrict the gRNAs discovered to only those for which the putative active site intersects the feature. There are additional options for filtering poly-T or poly-G tracks, restriction enzyme sites, and GC percentage. The user can also define a context window around the protospacer, which is a requirement for many downstream scoring methods. The default settings are for SpCas9 (NGG PAM, putative cleavage 4 bps upstream from the PAM) and Ruleset 3 scoring (30-mer target sequence centered on the protospacer).

### Building off-target index

Off-target scoring is performed by GuideScan2, and the wrapper script `index_genome.py` is used to generate the index from a fasta file. The wrapper script includes the option to subset the fasta in order to calculate the off-target effects against user-defined areas of interest (**Fig 2 C, D**).

### Scoring guide RNAs

Following gRNA identification, the output is passed to `score_guides.py,` which calculates the GuideScan2 off-target scoring and Ruleset 3 on-target scoring for each guide. Both scoring methods include multithreading, and the user can specify the number of gRNAs to process concurrently in order to avoid excessive memory usage on large sets (**Supp Fig 1B**). An arbitrary number of GuideScan2 indices can be passed, in which case each gRNA will be scored against each index.

If the user is interested in scoring Cas12a gRNAs or using an alternative scoring method for Cas9, the helper script `crisprscore.R` can be used. This script formats the input and applies an available scoring method from the crisprVerse Bioconductor package. Notably, any number of scoring methods can be applied and filtered over.

### Ranking guide RNAs

The final module, `rank_guides`, takes the scored guides as input and filters them according to user-defined criteria. Any number of scoring columns and minimum cut-offs can be used to filter out undesirable gRNAs. If the targets are genes represented by a GTF, the user can also specify the percentile range of the gene feature. For example, ‘`--percentile_range 0 50 --feature CDS`’ will filter out guides that are not within the first 50% of the coding sequence of any gene in the GTF. By default, min-max normalization is applied to scoring columns prior to standardizing scores before applying the final ranking.

### Model organism gRNA scoring

For the human genome sequence, GCA_000001405.15_GRCh38 was used with alternate haplotypes and mitochondrial chromosomes removed, and pseudoautosomal Y regions hard masked (converted to Ns). Genome sequences hosted at the UCSC Genome Browser were obtained for model organisms, and alternate haplotypes, mitochondrial DNA, and Y chromosomes were discarded. For Cas9 gRNAs, `generate_guides` was run with default settings to find all gRNAs that are expected to cleave in the CDS of any NCBI RefSeq gene annotation (curated and predicted gene models). `score_guides` with Ruleset 3 scoring with both tracrRNA options (Hsu2013, Chen2013) was run. For DeepHF scoring, both the “T7” and “U6” options were run, and the Cas9 enzyme was set to “WT” in both cases. For other scoring methods, default settings were used.

### Human cell line and mouse leukocyte consensus models

Transcript expression data was downloaded from the Encode consortium for each cell type using Encode’s uniform processing pipeline with Kallisto. Mapping is done against gencode.v29.annotation.gtf and gencode.vM21.annotation.gtf annotations for human and mouse, respectively. `preprocess_annotation` module was run in consensus mode with `--min` flag set to 0.1, 1, or 10.

### K562 transcription factor binding sites and off-target index

931 datasets were downloaded from the Encode K562 repository containing processed ChIP-Seq, ATAC-Seq, DNase-Seq, and ChIA-Pet in BED format. Hg19 datasets were removed, as were ChIP-Seq data indicating inactive genomic regions: H3K27me3 and H3K9me3. This left 886 datasets (Supplementary Table 13). A K562 active genome was constructed by passing these BED files, along with the comprehensive Gencode v45 annotation, to the `index_genome` module. The argument `--window 1000 1000` was used to expand the region around each active interval by 1000 bps in each direction to account for the local range of CRISPRi influence when calculating off-targets. For each transcription factor (TF) noted in Figure 2B, IDR thresholded peaks in BED format were downloaded from Encode, and each was run through the CRISPRware pipeline with default settings.

### Cast/C57 Hybrid Line

A C57BL/6 mouse was crossed to a Castaneous mouse to generate an F1 hybrid mouse line (Cast/C57). Legs were removed from a 12 week old F1 mouse and bone marrow was extracted as previously described (**PMID: 29761386**). Cells were grown in DMEM supplemented with 10% FCS, 5ml pen/strep (100X), 500ul ciprofloxacin (10 mg/ml), and 10% L929 supernatant (which contains macrophage colony-stimulating factor) with the replacement of culture medium every 2 to 3 d. On day 3, cells were infected with CreJ2 virus to induce immortalization as previously described (**PMID: 29761386**). After 3 months, when cells were capable of doubling without supplementation with L929, they were deemed fully immortalized (iBMDM-Cast/C57). The iBMDM-Cast/C57 were then lentivirally infected with the dCas9 construct that was constructed using Lenti-dCas9-KRAB-blast, addgene#89567. Cells were clonally selected for knockdown efficiency greater than 90%.

### Allele-specific targeting

Castaneous SNPs were downloaded from the Mouse Genome Database (PMID: 33231642) hosted at Mouse Genome Informatics (https://www.informatics.jax.org/). A Castaneous genome was generated using CRISPRware library script `replace_snps.py GRCm39.primary_assembly.genome.fa Cast.genome.fa Castaneous_SNPs.txt`. gRNAs were generated and scored through the standard pipeline against both the GRCm39 assembly (C57) and the Castaneous genome against TSS and CDS regions (gencode.vM34.annotation.gtf) on canonical chromosomes. Scored gRNAs were processed with a CRISPRware library script, with minimum hamming distance set to “1”, `allele_specific_guides.py C57_gRNAs Cast_gRNAs -d 1` This script produces three outputs: C57 specific protospacers and Castaneous specific protospacers which indicate a lost or gained PAM, and protospacers where the PAM is maintained but there is at least one SNP in the protospacer. Eipr1 was identified as a strong candidate for its baseline RNA expression, open chromatin at the TSS, the presence of a C57-specific PAM in the open chromatin, and documented SNPs in exon 8 for confirmation of expression change (amplicon-seq).

## Availability

CRISPRware code and documentation is maintained at https://github.com/ericmalekos/crisprware

## Acknowledgements

S. Carpenter is supported by R35GM137801 from NIGMS, and E.M. is supported by F31AI179201 from NIAID.

The authors would like to thank Prof. Angela Brooks for advice and guidance on the project.

## Author Contributions

E.M and S.C designed the research. E.M developed, tested, documented and implemented CRISPRware. C.M and S.C generated and tested the Cast/C57 Hybrid Line.

