## supplemental figures for "CRISPRware: an efficient method for contextual gRNA library design"

A

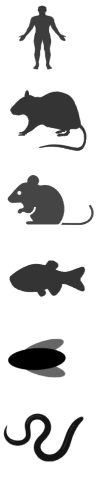

| | Coding genes | Coding transcripts | TTTV CDS protospacers | 5th to 65th % CDS | DeepCpf1 > 0.5 | gRNA/ gene | EnPAMGB > 0.5 | gRNA/ gene |
|---|---|---|---|---|---|---|---|---|
| **Hg38** | 20,627 | 139,092 | 1,084,770 | 630,252 | 420,578 | 12 | 496,119 | 14 |
| **Rn7** | 22,228 | 74,754 | 1,031,215 | 639,629 | 446,590 | 13 | 517,760 | 15 |
| **Mm39** | 22,864 | 98,005 | 1,056,344 | 656,290 | 457,223 | 13 | 529,667 | 15 |
| **DanRer11** | 26,424 | 57,093 | 1,608,055 | 771,550 | 543,521 | 15 | 607,824 | 16 |
| **Dm6** | 13,968 | 30,717 | 557,508 | 337,238 | 231,203 | 11 | 277,448 | 13 |
| **Ce11** | 20,093 | 28,146 | 1,144,629 | 709,498 | 451,266 | 19 | 582,607 | 24 |

B

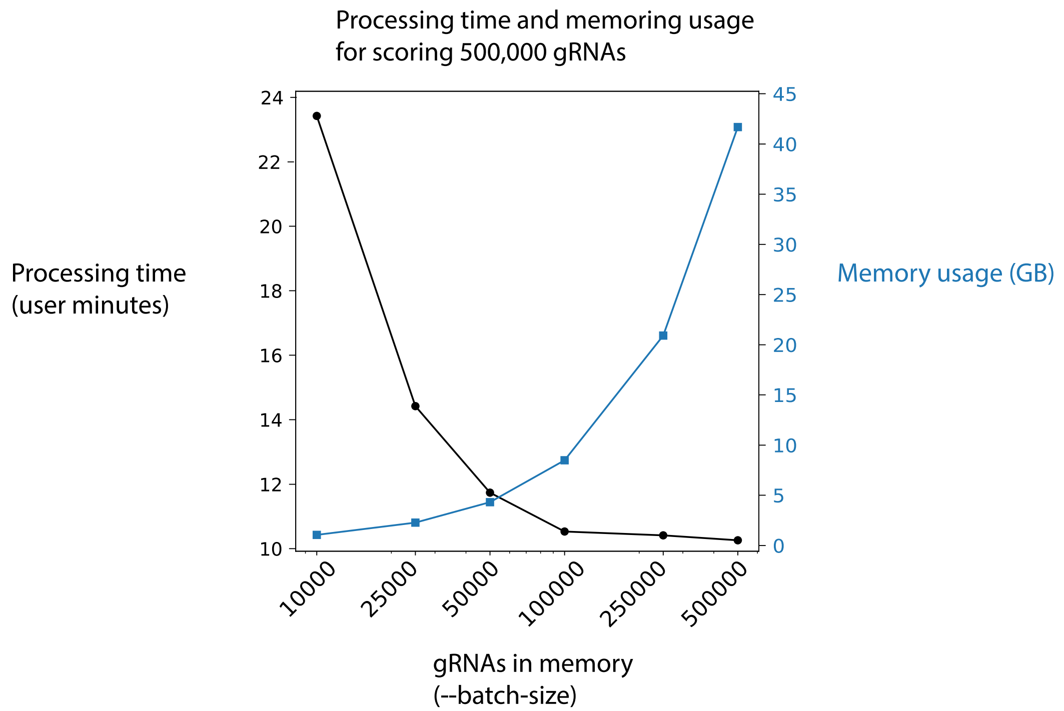

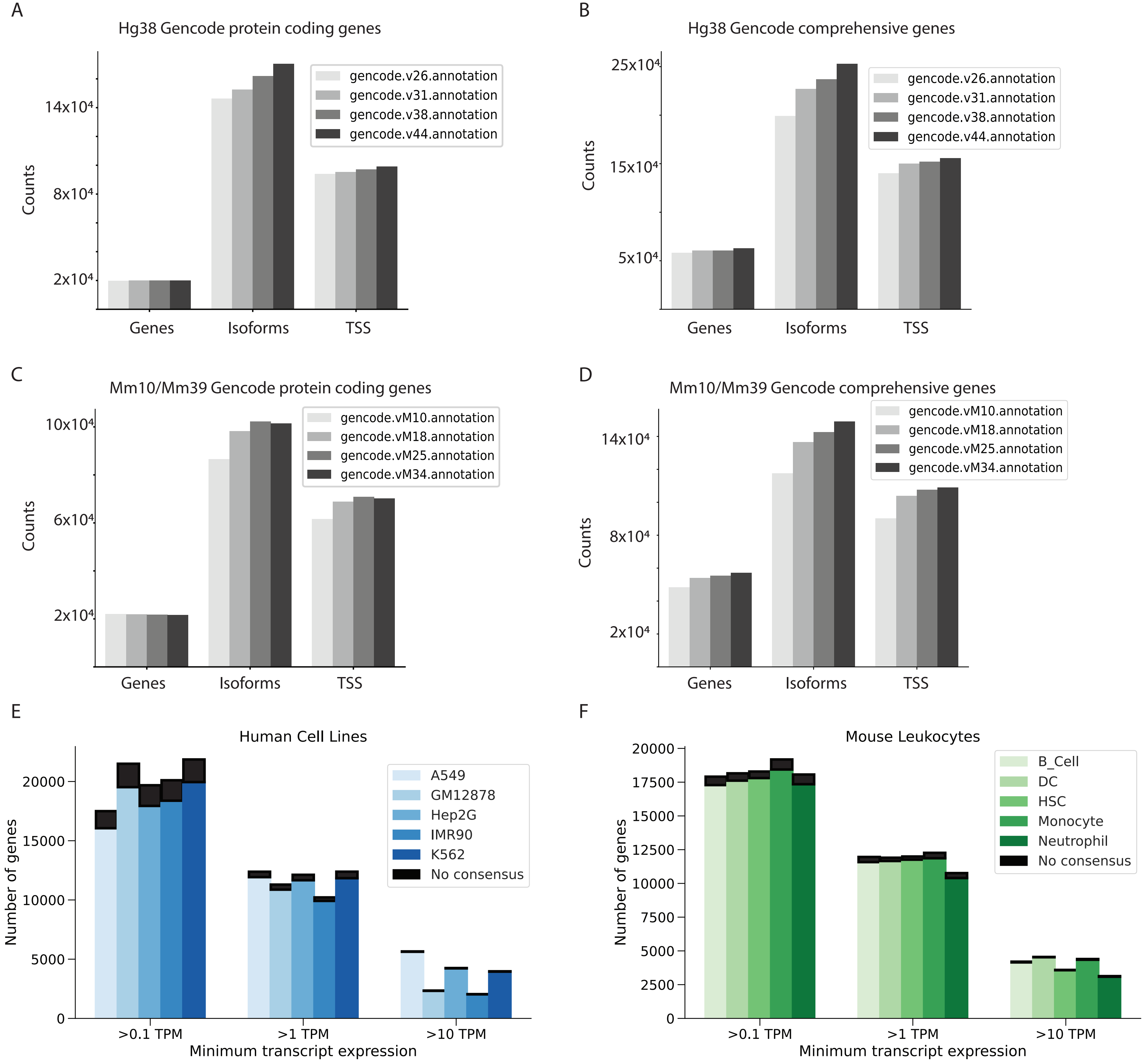
A
Hg38 Gencode protein coding genes
gencode.v26.annotation
gencode.v31.annotation
gencode.v38.annotation
gencode.v44.annotation
Counts
14x10⁴
8x10⁴
2x10⁴
Genes
Isoforms
TSS
B
Hg38 Gencode comprehensive genes
25x10⁴
15x10⁴
5x10⁴
C
Mm10/Mm39 Gencode protein coding genes
gencode.vM10.annotation
gencode.vM18.annotation
gencode.vM25.annotation
gencode.vM34.annotation
10x10⁴
6x10⁴
2x10⁴
D
Mm10/Mm39 Gencode comprehensive genes
14x10⁴
8x10⁴
2x10⁴
E
Human Cell Lines
A549
GM12878
Hep2G
IMR90
K562
No consensus
Number of genes
Minimum transcript expression
>0.1 TPM
>1 TPM
>10 TPM
F
Mouse Leukocytes
B_Cell
DC
HSC
Monocyte
Neutrophil
No consensus

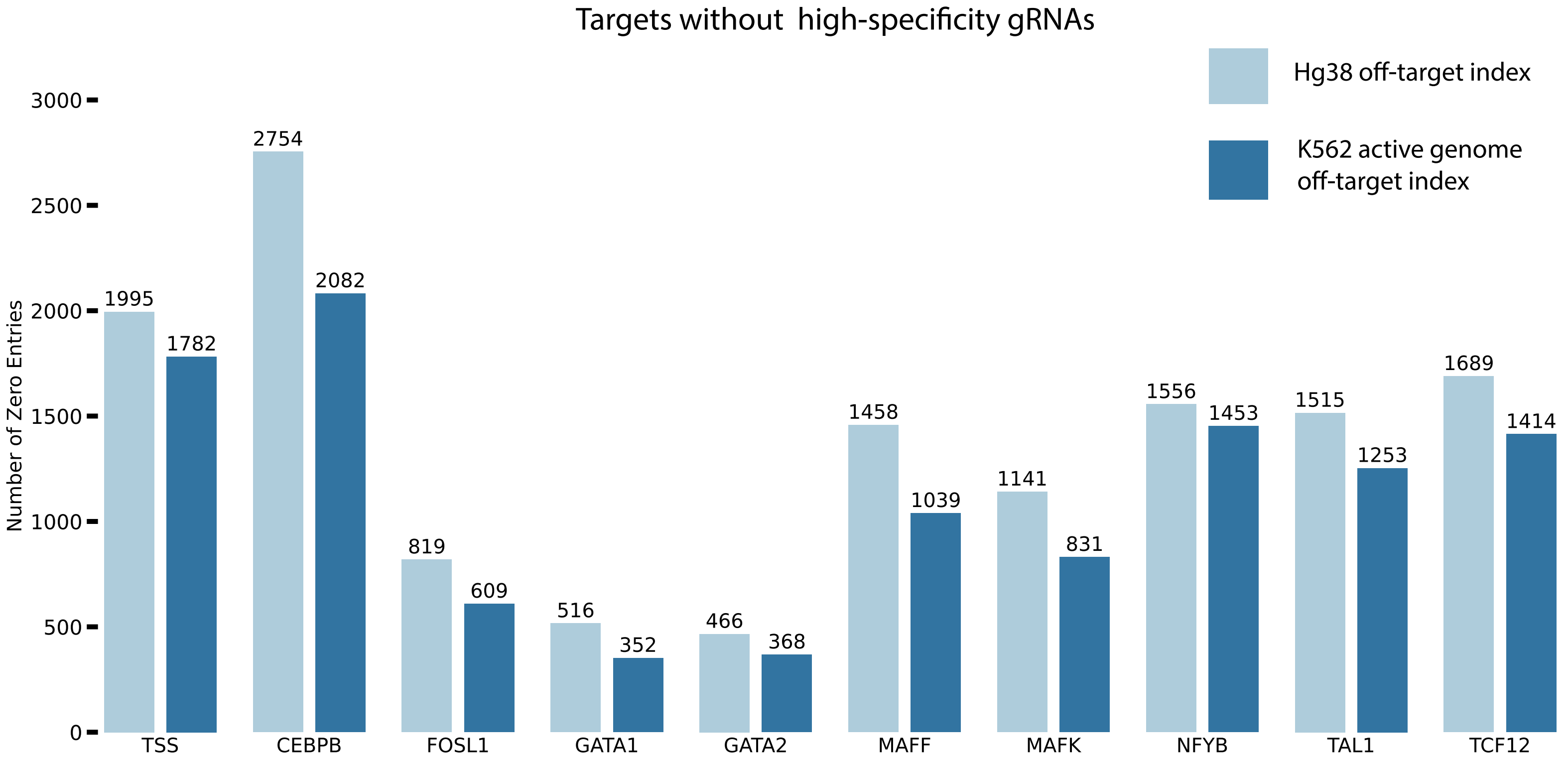
Targets without high-specificity gRNAs
Hg38 off-target index
K562 active genome off-target index
Number of Zero Entries
3000
2500
2000
1500
1000
500
0
1995
1782
2754
2082
819
609
516
352
466
368
1458
1039
1141
831
1556
1453
1515
1253
1689
1414
TSS
CEBPB
FOSL1
GATA1
GATA2
MAFF
MAFK
NFYB
TAL1
TCF12

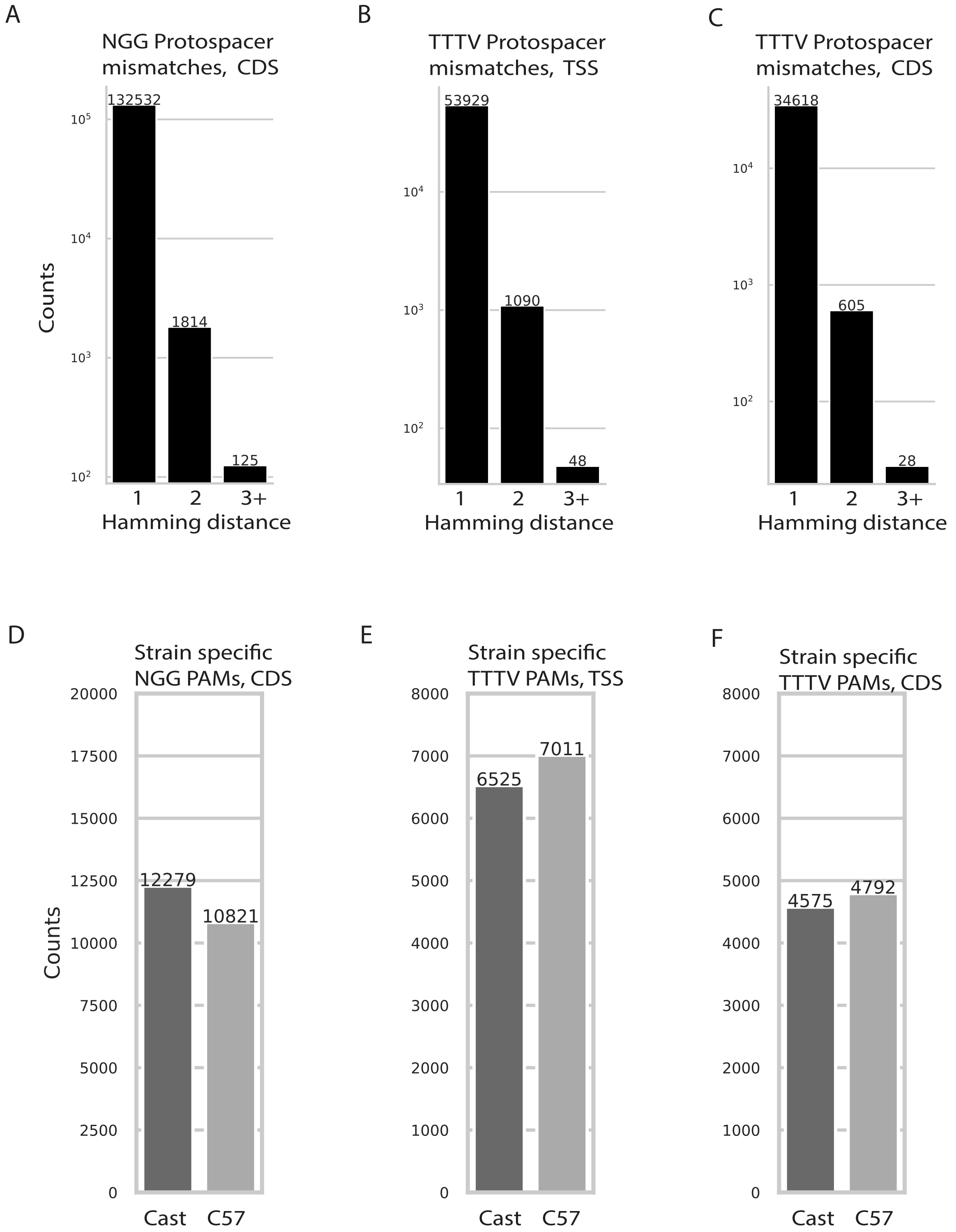
A
NGG Protospacer mismatches, CDS
132532
1814
125
$10^5$
$10^4$
$10^3$
$10^2$
Counts
1
2
3+
Hamming distance
B
TTTV Protospacer mismatches, TSS
53929
1090
48
$10^4$
$10^3$
$10^2$
1
2
3+
Hamming distance
C
TTTV Protospacer mismatches, CDS
34618
605
28
$10^4$
$10^3$
$10^2$
1
2
3+
Hamming distance
D
Strain specific NGG PAMs, CDS
20000
17500
15000
12500
10000
7500
5000
2500
0
Counts
12279
10821
Cast
C57
E
Strain specific TTTV PAMs, TSS
8000
7000
6000
5000
4000
3000
2000
1000
0
6525
7011
Cast
C57
F
Strain specific TTTV PAMs, CDS
8000
7000
6000
5000
4000
3000
2000
1000
0
4575
4792
Cast
C57
